# CrypToth: Cryptic pocket detection through mixed-solvent molecular dynamics simulations based topological data analysis

**DOI:** 10.1101/2024.07.10.602991

**Authors:** Jun Koseki, Chie Motono, Keisuke Yanagisawa, Genki Kudo, Ryunosuke Yoshino, Takatsugu Hirokawa, Kenichiro Imai

## Abstract

Some functional proteins undergo conformational changes to expose hidden binding sites when a binding molecule approaches their surface. Such binding sites are called cryptic sites and are important targets for drug discovery. However, it is still difficult to correctly predict cryptic sites. Therefore, we introduce a new method, CrypToth, for the precise identification of cryptic sites utilizing the persistent homology method. This method integrates topological data analysis and mixed-solvent molecular dynamics (MSMD) simulations. To identify hotspots corresponding to cryptic sites, we conducted MSMD simulations using six probes with different chemical properties: benzene, isopropanol, phenol, imidazole, acetonitrile, and ethylene glycol. Subsequently, we applied our topological data analysis method to rank hotspots based on the possibility of harboring cryptic sites. Evaluation of CrypToth using nine target proteins containing well-defined cryptic sites revealed its superior performance compared to recent machine-learning methods. As a result, in 7 out of 9 cases, hotspots associated with cryptic sites were ranked highest. CrypToth can explore hotspots on the protein surface favorable to ligand binding using MSMD simulations with six different probes and then identify hotspots corresponding to cryptic sites by assessing the protein’s conformational variability using the topological data analysis. This synergistic approach facilitates the prediction of cryptic sites with high accuracy.

## INTRODUCTION

The conformational diversity of protein, emerging from protein dynamics, results in multiple functional structures^1^. Conformational changes based on structural fluctuations or interactions with a small compound can create functional pockets not visible in ligand-free crystal structures; these are termed cryptic pockets^2^. Cryptic pockets, located away from active sites, and binding small compounds to such pockets could control protein function allostery. Allosteric modulators frequently target flexible regions, including cryptic pockets, because the mechanisms of protein allostery depend on the dynamic nature of proteins^2, 3^. Therefore, cryptic pockets represent valuable targets to expand the scope of drug discovery in terms that identification of the pockets is likely to lead to the discovery of allosteric modulators for inhibition or activation of protein function and drugs for proteins considered undruggable^4^. However, most cryptic pockets have been serendipitously discovered when both open and closed ligand-binding sites are observed in experimental protein structures^2, 5^. The deliberate identification of cryptic pockets is challenging, and the commonly used definition of cryptic pocket is ambiguous. For example, in developing a dataset of 93 proteins for a cryptic site prediction tool, Cryptosite^6^, a site is classified as cryptic if it remains undetected by FPocket^7^ or ConCavity^8^ when comparing a bound structure to an unbound structure. However, approximately half of proteins possess at least one unbound structure capable of binding a ligand without causing clashes with the protein^9^.

Given the dynamic nature of proteins, molecular dynamics (MD) simulations offer a promising approach for exploring cryptic pockets^10^. Simply identifying a cryptic pocket is insufficient for drug target discovery; it must also be ligandable. Mixed-solvent MD (MSMD) simulations involve small probes (chemical fragments) such as isopropanol and benzene mixed with water molecules. MSMD simulations enable the observation of protein flexibility and the mapping of “hotspots” on protein surfaces, which are identified by the probe with higher occupancies on the protein surface during the MD simulation. Thus, MSMD simulations enhance the sampling of cryptic pocket formations concurrently by estimating the pocket location based on mapping hotspots. Various methods have been utilized to explore cryptic pockets^11–15^; however, several hotspots are generally observed on a protein surface. Therefore, one of the goals of the MSMD protocol is to accurately identify a hotspot corresponding to the cryptic site location among all hotspots. Kimura et al. selected eight proteins from the Cryptosite dataset^6^ and successfully mapped six of the eight cryptic pockets within the top three hotspot ranks. They conducted 100 ns MD simulations on each system using isopropanol, resorcinol, or acetic acid at 5% concentrations, ranking the sites based on free energy computed from probe occupancy^11^. Schmidt et al. performed 100–500 ns MSMD simulations with 10% phenol on seven proteins to identify pocket cores overlapping with proposed cryptic pockets in the proteins. They managed to detect these sites on six of the seven proteins using the top three hotspot ranks^12^. Smith and Carlson selected 12 proteins exhibiting diverse conformational changes, and their dataset did not overlap with that of Kimura et al. They conducted 100 ns conventional mixed-solvent and accelerated mixed-solvent MD simulations, identifying cryptic sites in seven proteins using five different probes and hotspot mapping^15^. These studies demonstrate the challenge of identifying hotspots corresponding to cryptic pockets solely using probe-occupancy-based hotspot ranking. Furthermore, observing the exposure of cryptic pockets poses a significant challenge^11, 12, 14^. The chemical properties of the probe play a crucial role in identifying cryptic pockets, leading to the use of various probe molecules with different chemical properties, such as benzene, isopropanol, phenol, imidazole, acetonitrile, and ethylene glycol^15^. However, none of these probe molecules have been reported as effective.

Therefore, identifying hotspots corresponding to cryptic sites is challenging. Considering that structural changes are a key factor in cryptic site formation, hotspots should be evaluated based on the dynamic features of proteins around cryptic sites. In this study, we developed CrypToth (Figure 1), a method for the detection of cryptic pockets. This method combines the exploration of hotspots on ligandable protein surfaces using MSMD simulations with six probe molecules and the evaluation of protein structural variability using the previously developed topological data analysis method DAIS (Dynamical Analysis of Interaction and Structural changes)^16^. The efficacy of CrypToth was validated using nine proteins containing cryptic sites undetectable by conventional pocket search methods. In addition, we compared the capability of detecting cryptic sites between CrypToth and the PocketMiner method^17^, a machine-learning-based prediction method.

**Figure 1.**
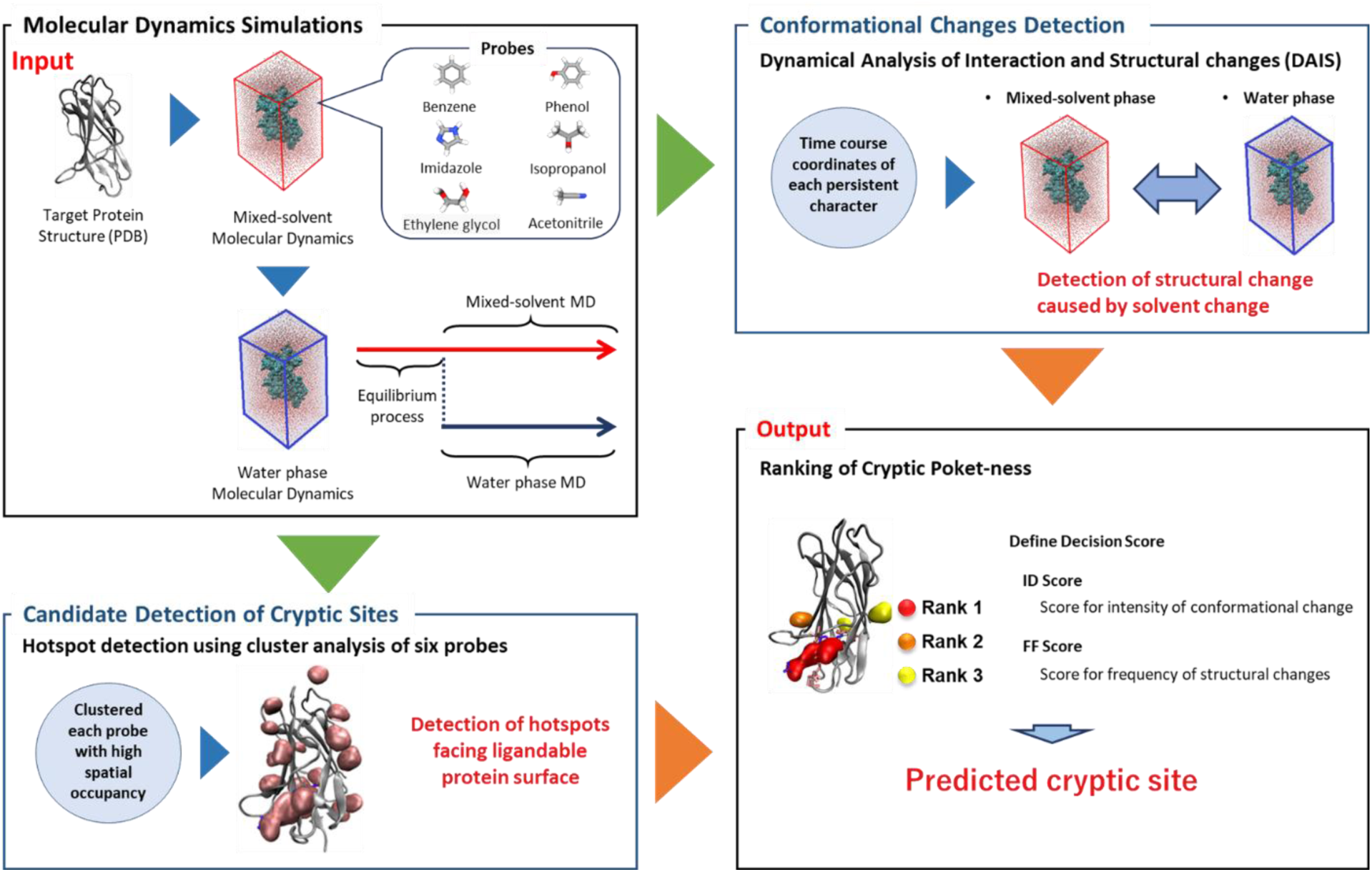
Schematic of the cryptic site detection flow (CrypToth) using MSMD simulations and topological data analysis via the DAIS method.

## METHODS

### Choice of Proteins with Cryptic Pockets

To evaluate our method, we selected 66 candidates (pairs of apo and holo structure) with cryptic sites from previous studies^11–13, 15, 17^. In this study, we defined a cryptic site as the one that clashes with the ligand molecule in apo structure when apo structure is superimposed on holo structure, and also is undetectable through pocket search against apo structures. Based on this definition, nine target proteins were selected from the candidate proteins (Table 1). Detection of the clashing residue is as follows. We determined clashes between residues and the ligand by assessing the distance between the heavy atoms of the ligand molecule in the holo structure and those of the protein residue in the apo structure, considering a distance of less than 3.5 Å as indicative of a clash. When multiple pairs of apo and holo structures were available, we selected holo structures binding to different ligand as representative holo structures and then the ligand binding site are defined as the union of the residues which Meanwhile, we selected multiple representative apo structures through clustering analysis based on structural comparisons of the ligand binding site. Subsequently, we identified residues clashing with each ligand based on the ratio of clashes in the representative apo structures (Tables S1-5); while the multiple ligand binding sites of the representative apo structures are superimposed to that of a holo structure, a residue of the apo structures clashing with the ligand with a ratio of ≥ 0.8 was considered to clash with the ligand. The clustering of apo structures was conducted using the “clustering of conformers function” in Schrödinger Suite 2020-4^20^, with the number of clusters determined by the Kelly criterion. The centroid structure within each cluster was designated as the representative structure. Additionally, we utilized SiteMap, a pocket search tool within the Schrödinger Suite 2020-4^18^, to identify apo proteins with undetectable cryptic sites.

**Table 1.** Proteins with cryptic sites are selected from the present work. The name of each protein, the PDB IDs (apo and holo), and the residues that inhibit ligand binding are shown.

| Protein Name | PDB ID |  | Clashed residues* |
| --- | --- | --- | --- |
|  | apo | holo |  |
| TIE-2 | 1FVR | 2OO8 | ILE830, GLY831, GLU832, ASN834, PHE835, GLY836, VAL838, LYS855, ILE902, GLU903, ALA905, HIS962, ILE980, ALA981, ASP982, PHE983, GLY984, LEU985 |
| Exodeoxyribonuclease I | 1FXX | 3HL8 | TRP245, ALA312, ARG327, CYS330, LEU331, ARG338 |
| Beta-lactamase TEM | 1JWP | 1PZO | PRO217, LEU218, LEU219, SER221, ALA222 |
| NPC intracellular cholesterol transporter 2 | 1NEP | 2HKA | TYR55, VAL57, VAL83, PHE85, LEU113, PRO114, VAL115, LYS116, GLU118, TYR119, PRO120, ILE145 |
| Androgen receptor | 2AM9 | 2PIQ | VAL717, LYS721, VAL731, MET735, ILE738, GLN739 |
| Dihydrofolate reductase type 1 from Tn4003 | 2W9T | 2W9S | ASN19, GLN20, LEU21, PRO22, TYR23, ASP28, LEU29, ILE32, PHE93 |
| Ferulic acid decarboxylase | 3NX1 | 3NX2 | TYR21, ANS23, GLY24, TRP25, TYR27, PHE95, ILE1312 |
| Fascin | 3P53 | 6I11 | PHE14, LEU16, ALA58, ALA59, VAL60, LEU92, ILE93, VAL94, ALA95, TRP101, LEU103, GLU215, PHE216, ARG217, ARG224, TYR230 |
| Anti-methotrexate CDR1-4 Graft VHH | 3QXW | 3QXV | GLN3, VAL4, ALA26, ARG28, SER30, SER31, ARG32, MET36, ARG74, ASN76, TYR79, ALA100 |

\*Clashed residues located near hotspots matching the cryptic sites are shown. Residue numbers were based on sequence numbers obtained from UniProt<sup>19</sup>. For the Anti-methotrexate CDR1-4 Graft VHH, the residue numbers were based on the sequence numbers in PDB (3QXW).

### Development of CrypToth

CrypToth conducts hotspot detection and identifies conformational changes using MSMD and DAIS, respectively (Figure 1). Here, we describe the methods used for hotspot detection and conformational change identification.

### MSMD simulations

To conduct MSMD simulations, we utilized the Protein Preparation Wizard in the Schrodinger suite 2020-4^18^ to complete the absent loops and side chains for all target proteins. During this procedure, the N- and C-termini were capped with N-methyl amide and acetyl groups, respectively. Subsequently, non-protein molecules were eliminated, and structural optimization was performed using the OPLS3e force field^19^. The RESP charges for the six probe molecules used in this study (benzene, isopropanol, phenol, imidazole, acetonitrile, and ethylene glycol) were calculated at the HF/6-31G* level using the Gaussian 16 software package^20^.

The MSMD simulation protocol described below was conducted using EXPRORER_MSMD^22, 23^. To generate the MSMD system, PACKMOL 18.169^21^ was used to randomly place probe molecules around each protein, equilibrated in a water-phase environment at 0.25 M, followed by the addition of water molecules. The AMBER ff14SB^22^, GAFF2^23^, and TIP3P^24^ force fields were used for the proteins, probe molecules, and water molecules, respectively. Additionally, a Lennard-Jones force field term was introduced (ε_*LJ*_ = 10^−6^ kcal/mol; Rmin = 20 Å) solely between the center of the probes to prevent aggregation.

After the energy minimization of the above MSMD systems, NVT (canonical ensemble) MD simulations with harmonic position restraints on the solute heavy atoms (force constant, 10 kcal/mol/Å^2^) were performed to gradually heat the system up to 300 K over 200 ps. Subsequently, NPT (isothermal–isobaric ensemble) MD simulations of 800 ps at 300 K and 10^5^ Pa were performed to gradually remove the position restraints. The P-LINCS algorithm^25^ was utilized to constrain all bond lengths, including hydrogen atoms, with a time step (⊿t) of 2 fs. In this study, 20 runs of 40 ns MSMD were executed for each system, and the trajectories of the final 20 ns were analyzed for hotspot detection. Trajectories of 5 ns (20-25 ns) were specifically used for DAIS analysis. These simulations were conducted using GROMACS 2021.5^26^.

### Detection of hotspots corresponding to cryptic sites

To identify hotspots in each MSMD simulation, we initially determined the occupancy of probe heavy atoms for each probe in a voxel (1 Å × 1 Å × 1 Å grid voxel). These computations utilized the PMAP^27^. In this study, owing to variations in the number of heavy atoms across the probe molecules used, probe occupancy was normalized by dividing the occupancy value by the number of heavy atoms in each probe molecule. Subsequently, probe occupancies exceeding 0.0004 were selected and clustered using the DBSCAN algorithm^28^ to identify clusters (DBSCAN parameters: ε_*DBSCAN*_ = 3.0; min_samples = 7). These identified clusters correspond to regions in the MSMD simulation where probe molecules tended to stay on the protein surface.

After extracting clusters for each probe, clusters with over 20% overlap were integrated into a single hotspot. This integration of clusters containing multiple probe molecules incorporates the properties of the binding ligand into the hotspots. A hotspot was defined as a “cryptic site” if more than 50% of its voxels were within 3.5 Å of any residue atoms that clash with the ligand molecule, and moreover any voxels were within 4.5 Å of any ligand atom.

### Evaluation of protein conformational changes based on structural differences in aqueous and mixed-solvent phases

The DAIS method was used to rank the possibilities of cryptic sites as hotspots. This method is based on persistent homology and can extract significant structural changes, no matter how small, between the target and reference structures. To prepare for structural change analyses using the DAIS method, all solvent molecules were removed from structural conformations that had been fully thermally equilibrated in a mixed-solvent environment with each probe. Subsequently, structural sampling was conducted for 5 ns in an aqueous environment under identical conditions to the MSMD simulation. For the conformational change analysis with DAIS, 500 snapshots of structural conformations from the mixed solvent and water phases were used. These analyses used five trajectories for each probe.

To evaluate the potential of cryptic sites as hotspots, we analyzed the differences between the protein structures derived from MD simulations in mixed solvent and water-phase environments using the DAIS method. Two scoring metrics, the absolute ID score and FF score, were used to determine the likelihood of a site matching a hotspot being a cryptic site. The ID score is the original score computed through the DAIS method^16^. It quantifies the structural variance between the reference and target structures. This score reflects the extent of structural alteration, as defined by the equation:

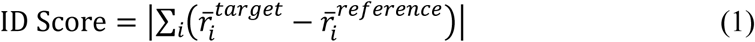

where *r^-^*_*i*_^*target*^ and *r^-^*_*i*_^*reference*^ correspond to the magnitude of the variability of the substructure captured by the DAIS method. The score can be positive or negative, indicating which of the compared structures is more mobile. However, absolute values were used in this study to focus solely on the magnitude of the structural change, and those with a score of 5.0 or greater were selected. Additionally, a score was adopted that averaged the ID scores possessed by the residues forming the topology, which were identified as structurally different using the DAIS method. The FF score is a new score introduced in this study and represents the feature-point frequency. This score is a count of the number of residues around each candidate hotspot with Cα atoms forming the topology determined to have a structural difference by the DAIS method, as shown in the following equation:

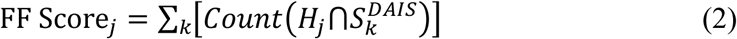

where *H*_*j*_ refers to the set of Cα atoms from the residues around hotspot *j*, identified through MSMD simulations, while *S*^*DAIS*^_K_ denotes the feature point *k*, which consists of the set of Cα atoms identified using the DAIS method. These scores were calculated for each MSMD calculation and subsequently used to rank the potential of the candidate to be a cryptic site. The ID and FF scores were calculated from the five trajectories for each hotspot using each probe. In this study, we omitted cases in which the scores calculated from each trajectory deviated significantly with the goal of making highly accurate predictions from a small number of trials. Specifically, trials in which the score deviated more than the standard deviation from the mean were excluded as outliers, and the mean score was calculated for the remaining trials.

## RESULTS & DISCUSSION

### Structural characteristics of the target proteins

Nine proteins containing cryptic sites were used for the evaluation of CrypToth (Table 1, see METHODS). For all of these proteins, both the apo (unbound) protein structures and holo (bound) protein structures with the ligand molecule have been reported through X-ray crystallography. In addition, considering a practical case, we selected proteins whose cryptic sites in the apo structures could not be identified using the SiteMap method^29, 30^, which is one of the binding pocket search methods. As shown in Figures 2 and S1, the binding ligands in the holo state clashed with the residues surrounding those in the apo state when the apo and holo structures were superimposed. Purple bars and pink van der Waals spheres represent the ligand and colliding residues in the apo state, respectively. The residue groups that collided with the ligand in each apo structure are listed in Table 1. In most proteins, such as Exodeoxyribonuclease I (1FXX) and the androgen receptor ligand-binding domain (2AM9) (Figures 2A, B), the cryptic site is generated by changing the orientation of the side chain of residues within the ligand-binding region, although it is generally difficult to observe such change of side chain for opening of cryptic sites from apo structure using conventional MD simulation. Notably, this occurs without significant visible changes in the backbone structure. Conversely, proteins, such as anti-methotrexate CDR1-4 Graft VHH (3QXW) (Figure 2C) undergo substantial conformational change in specific regions to create binding sites.

**Figure 2.**
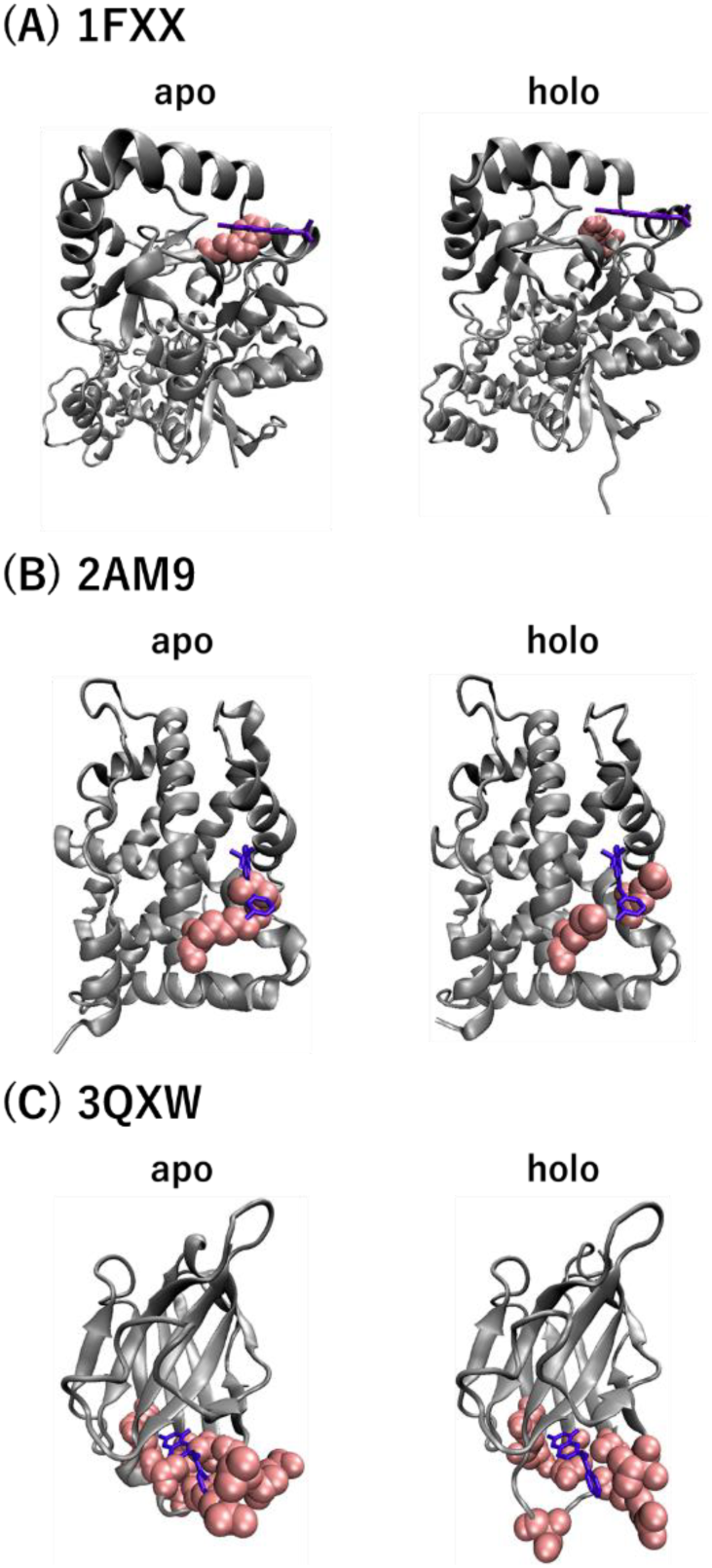
Representative conformational collision states between each ligand and protein at the cryptic site. The ligand positions in the holo state are displayed on the superimposed apo structure, illustrating the overlap between the ligand and amino acids. The purple bars and pink van der Waals spheres indicate the binding ligand and the colliding residues in the apo structure, respectively.

### Hotspot identification using MSMD simulations

To detect the possible cryptic sites in each target protein, MSMD simulations were performed using six probe molecules, including benzene, isopropanol, phenol, imidazole, acetonitrile, and ethylene glycol. In the MSMD trajectories, after sufficient equilibration, hotspots were detected based on the congregative occupancies of the six probe molecules on the protein surfaces (Figure S2, see METHODS). As shown in Figure S2, hotspots were observed at different locations on the surface of each protein. Except for beta-lactamase TEM (1JWP), hotspots located around the cryptic site were observed in all proteins. Although sufficient structural changes to open the cryptic sites were not observed during our MSMD simulation, hotspots were located on the cryptic sites. These observations confirmed that hotspot detection using MSMD simulations is useful for exploring cryptic sites.

### Discriminating hotspots corresponding to cryptic sites through structural change analysis

As shown in Figure S2, hotspots were observed at different locations on the surface of each protein. It is difficult to discriminate hotspots matching cryptic sites among all hotspots using hotspot ranking based only on probe occupancy. The structural changes induced by ligand binding are the essence of the cryptic site. Thus, CrypToth incorporated DAIS^16^ to estimate the conformational variability that emerges from interactions with chemical probes by comparing the MD trajectories in a water-phase environment and in a mixed solvent environment. From the DAIS results, the “averaged ID score” and “FF score” were calculated for the residues located within 3.5 Å of a hotspot. Between the target (mixed-solvent phase) and reference structures (water phase), the ID score indicates the magnitude of the structural change observed, whereas the FF score shows the observed frequency of the structural change.

Tables S6 and S7 present the results of hotspot ranking by ID and FF scores, respectively, obtained from the MSMD simulation with each probe. When using the ID score for ranking, no probes could detect the hotspot corresponding to the cryptic site within the top three for all target proteins. However, ranking of the FF score based on the MSMD simulation with phenol yielded significantly better results for all target proteins. Notably, hotspots corresponding to cryptic sites ranked first among the seven target proteins, except for the ligand-binding domain of the androgen receptor (2AM9). Interestingly, the hotspot corresponding to the cryptic site was ranked second in 2AM9. The ranking accuracy was higher with the FF score than with the ID score, indicating that the frequency of structural change occurrence is more essential for detecting cryptic sites than the magnitude of structural change.

To evaluate CrypToth, we conducted a performance comparison with PocketMiner^17^, a state-of-the-art graph neural network-based cryptic site prediction method. Based on the results, we selected regions consisting of residues with a prediction score of 0.7 or higher, following the threshold for cryptic site prediction described in the PocketMiner study^17^. The ability to detect cryptic sites was evaluated by counting the number of residues around a hotspot, which corresponded to the FF score. As shown in Table 2, CrypToth outperformed PocketMiner in terms of ranking accuracy. In the case of PocketMiner, the hotspot corresponding to the cryptic site was ranked within the top three in five target proteins (ranked first in only two targets). In contrast, CrypToth with phenol ranked the hotspot corresponding to the cryptic site first among the seven target proteins. Furthermore, the cryptic site hotspot was ranked second in the remaining target. To provide a more visual understanding of the results when ranking using each score (ID, FF, and PocketMiner scores), the top three hotspots are displayed for each structure (Figure 3). Compared with PocketMiner, the FF score ranking detected cryptic sites as rank 1 hotspots in almost all cases. In Figure S3, projection views are shown when using the results of the phenol MSMD simulations. In this figure, the top five ID and FF scores are mapped onto the protein structures with color gradation. In addition, each hotspot detected at rank 1 is highlighted. It is noteworthy that the ID and FF scores are not absolute values that can be directly compared across different proteins; therefore, their evaluation cannot rely solely on color representation.

**Figure 3.**
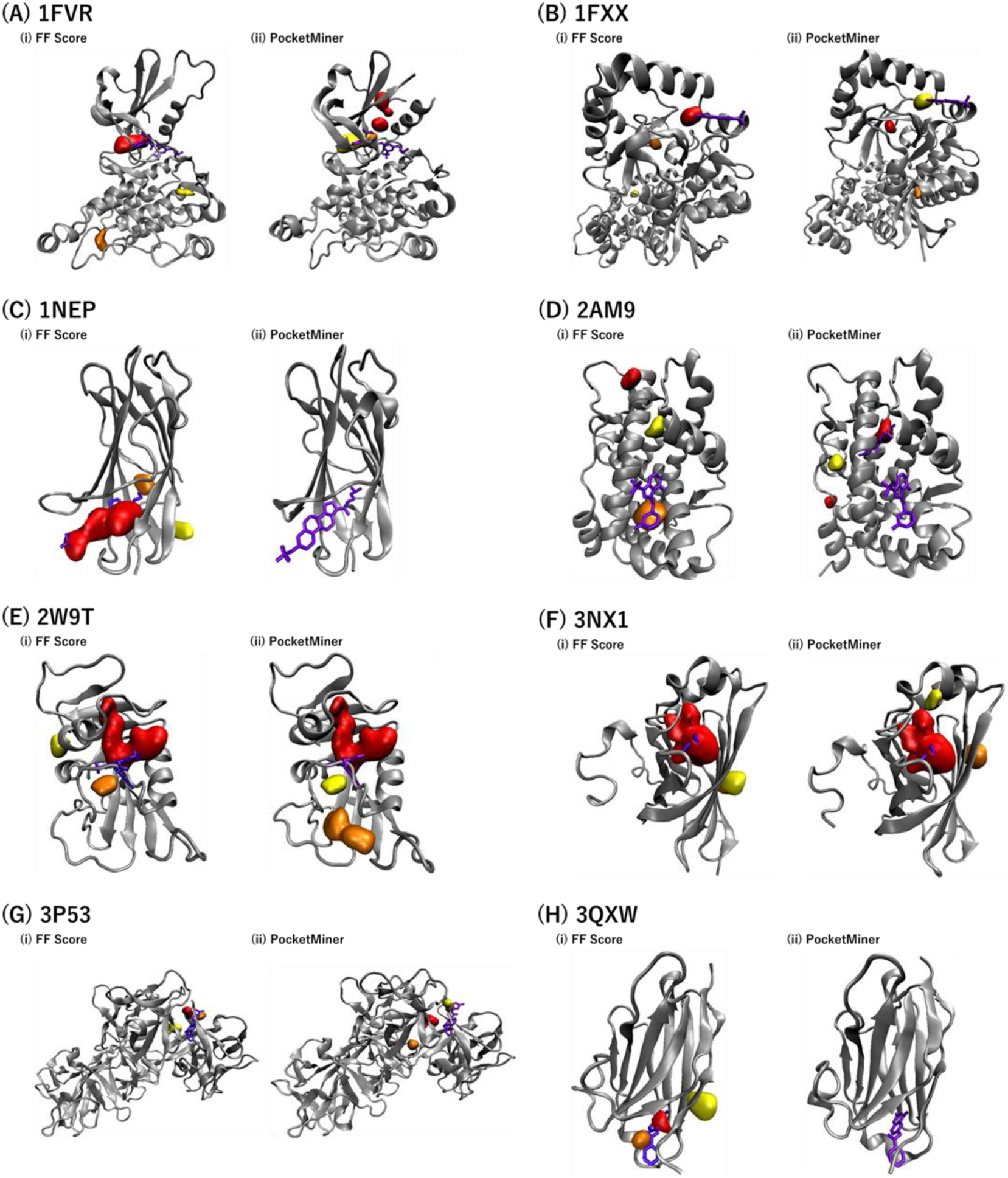
Ranking hotspots based on the estimation of the “structural changeability” of the protein. The cartoon representation in silver illustrates the protein backbone of the apo structure, while the purple stick representation indicates the ligand binding position in the holo structure. The pink stick representation shows the cryptic site residues that clash with the ligand. The hotspot most likely to be a cryptic site (rank 1) is shown in red, followed by rank 2 in orange and rank 3 in yellow.

**Table 2.** Comparison of hotspot rankings for cryptic sites between CrypToth and PocketMiner, with rankings in CrypToth based on the FF score calculated using MSMD simulations with phenol.

|  | CrypToth<br>(Phenol) | PocketMiner | Number of Hotspots |
| --- | --- | --- | --- |
| 1FVR | 1 | 2 | / 21 |
| 1FXX | 1 | 3 | / 6 |
| 1NEP | 1 | Not Detected | / 16 |
| 2AM9 | 2 | 11 | / 18 |
| 2W9T | 1 | 1 | / 16 |
| 3NX1 | 1 | 1 | / 16 |
| 3P53 | 1 | 3 | / 7 |
| 3QXW | 1 | Not Detected | / 12 |

Table 3 shows the relationship between the probe combination comprising the hotspot corresponding to cryptic sites and the probe yielding the best results in cryptic site detection. The FF score, calculated from MSMD simulations with phenol, demonstrated the most accurate detection of cryptic sites in nearly all cases. However, phenol was not included in all hotspots corresponding to cryptic sites, suggesting that the ligand binding space of a cryptic site may not solely result from the interaction between probe molecules and protein surface near the site. Instead, it shows the possibility that conformational changes propagate similarly to allosteric effects due to the interaction with probes on the protein surface far from the site. The CrypToth approach, which combines MSMD and DAIS methods, proves highly effective in identifying cryptic sites for two main reasons. First, MSMD can consider the ligandable surface by utilizing various probe molecules with different chemical properties. Second, DAIS can consider the structural change of the protein.

**Table 3.** Probes associated with hotspots corresponding to cryptic sites and optimal probes for cryptic site detection via DAIS analysis. When multiple hotspots correspond to a cryptic site, those are shown in different rows.

| PDB ID<br>(apo) | Probes consisting of Hotspot<br>matching Cryptic site | Optimal probe<br>in cryptic site detection |
| --- | --- | --- |
| 1FVR | Benzene, Isopropanol, Phenol, Imidazole,<br>Acetonitrile | Phenol, Imidazole |
|  | Ethylene glycol |  |
| 1FXX | Benzene, Acetonitrile | Phenol |
| 1JWP | No hotspot matching cryptic site |  |
| 1NEP | Benzene, Isopropanol, Phenol, Imidazole,<br>Acetonitrile | Phenol, Imidazole |
|  | Benzene |  |
| 2AM9 | Benzene, Isopropanol, Phenol | Phenol |
| 2W9T | Imidazole | Phenol, Imidazole,<br>Acetonitrile, Ethylene<br>glycol |
|  | Benzene, Isopropanol, Phenol, Imidazole,<br>Acetonitrile | Benzene, Isopropanol,<br>Phenol, Imidazole,<br>Acetonitrile, Ethylene<br>glycol |
| 3NX1 | Benzene, Isopropanol, Phenol, Imidazole,<br>Acetonitrile, Ethylene glycol | Benzene, Isopropanol,<br>Phenol, Imidazole |
| 3P53 | Acetonitrile | Phenol |
| 3QXW | Acetonitrile | Benzene, Phenol,<br>Imidazole, Ethylene glycol |
|  | Acetonitrile, Ethylene glycol |  |
|  | Acetonitrile |  |
|  | Benzene |  |

## CONCLUSIONS

We propose a new method, CrypToth, for detecting cryptic sites by combining the identification of ligandable surface using MSMD simulations and estimating structural changeability using DAIS. MSMD simulations with probe molecules of varying chemical properties facilitated the observation of the occupancy time at the interaction point for each molecular property. This approach is similar to identifying pharmacophore points in fragment-based drug discovery methods, where ligandability are considered in our hotspot calculations based on the occupancy of multiple fragments. Additionally, DAIS identifies structural differences between the target and reference structures, identifying regions where significant structural changes may occur in the future within a short simulation time. This capability is valuable for predicting cryptic sites, as it captures the structural changes when a region returns to its original stable configuration in an aqueous solution following the removal of the probe molecule from its equilibrated state in a mixed solvent. CrypToth maximizes these advantages, surpassing current machine learning methods. Future machine-learning-based forecasting models can leverage the features derived from this approach.

## ASSOCIATED CONTENT

### Data Availability

Initial 3D structures of the proteins were downloaded from the Protein Data Bank (PDB). Schrodinger suite 2020-4 was used for protein preparation. We used AmberTools21 and Gaussian 16 Rev B.01 for probes preparation. PACKMOL 18.169 and AmberTools18 were used to prepare the MSMD system. GROMACS 2021.5 was used as the MD engine. VMD 1.9.3 was used for visualization. The protocols of the MSMD simulation and DAIS in this study are available from https://github.com/keisuke-yanagisawa/exprorer_msmd and https://github.com/jkoseki/DAIS.

### Supporting Information

Supporting Information is available. Supporting Information: DOCX. Collision status of each ligand and protein at the cryptic site. (Figure S1) the hotspots obtained from MSMD simulations. (Figure S2); The projection views to compare three score values. (Figure S3) DAIS analysis results for 1JWP with a loose threshold. (Figure S4). Detection of the residues crashing with the ligands in each validation protein are shown in (Tables S1 to S5) and the rank order of scores obtained using DAIS methods for cryptic hotspot sites (Tables S6 and S7), respectively.

## AUTHOR INFORMATION

### Author Contributions

JK, CM, and KI designed the study and performed program creation and variation analyses. JK, CM, and KI discussed improvements in the research. CM and KI selected the target molecules with cryptic sites. CM, KY, and GK performed the mixed-solvent molecular dynamics simulations. JK analyzed the structural changes using DAIS. JK, CM, and KI wrote the manuscript. All the authors have read and approved the manuscript.

### Notes

The authors declare no competing interests.

## ACKNOWLEDGMENT

Computational experiments were partially performed using the Cygnus system at the CCS, University of Tsukuba.

## ABBREVIATIONS

MSMD: mixed-solvent molecular dynamics

DAIS: dynamic analysis of interactions and structural changes.

RESP: restrained electrostatic Potential

GAFF2: General AMBER Force Field 2.

