## Supplementary material for "CrypToth: Cryptic pocket detection through mixed-solvent molecular dynamics simulations based topological data analysis": Figure S1-S3 and Table S1-S7

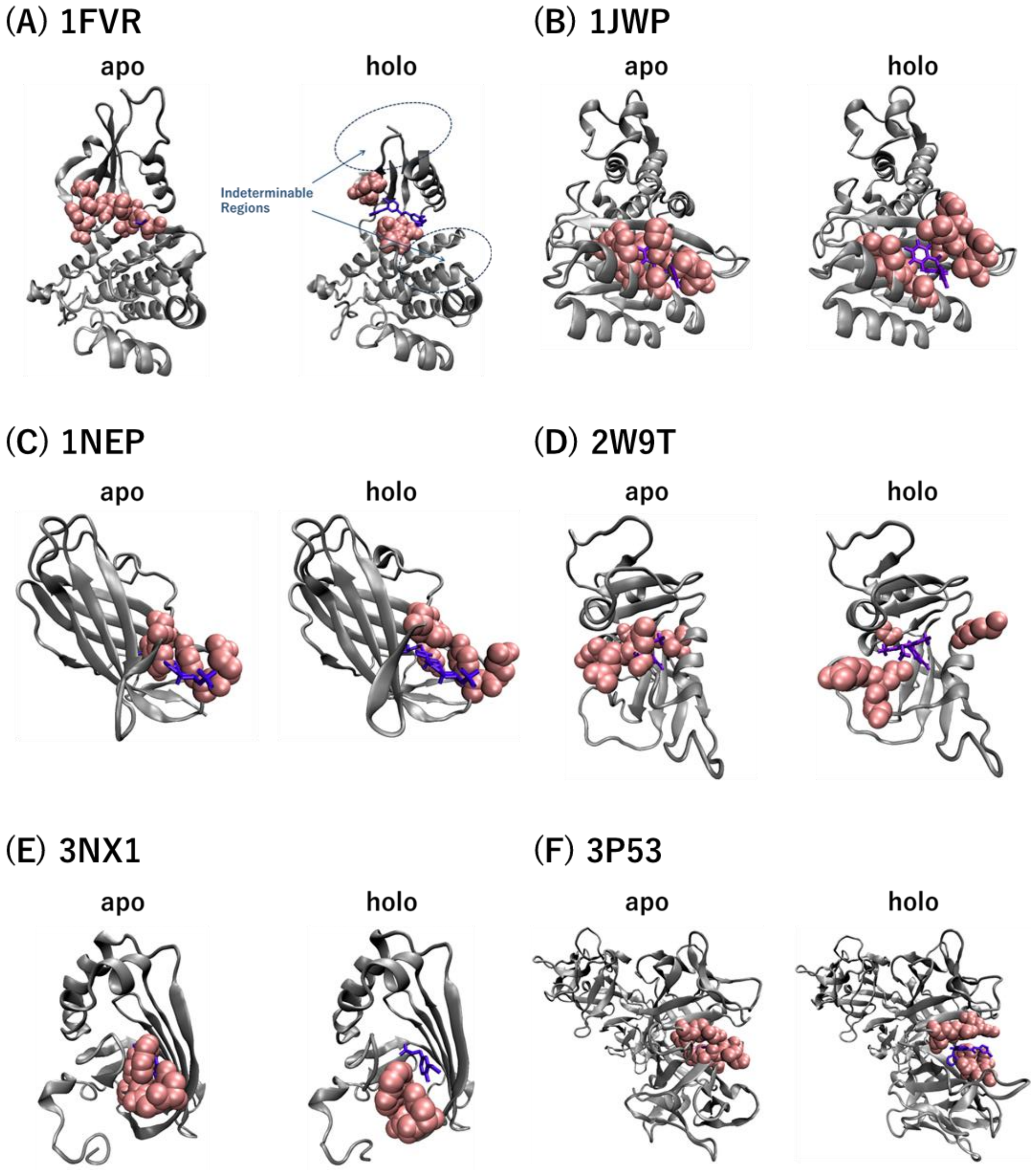


**Figure S1.** The collision states between each ligand and protein at the cryptic site. The ligand positions in the holo state are displayed on the superimposed apo structure, illustrating the overlap between the ligand and amino acids. The purple bars and pink van der Waals spheres indicate the binding ligand and the colliding residues in the apo structure, respectively.

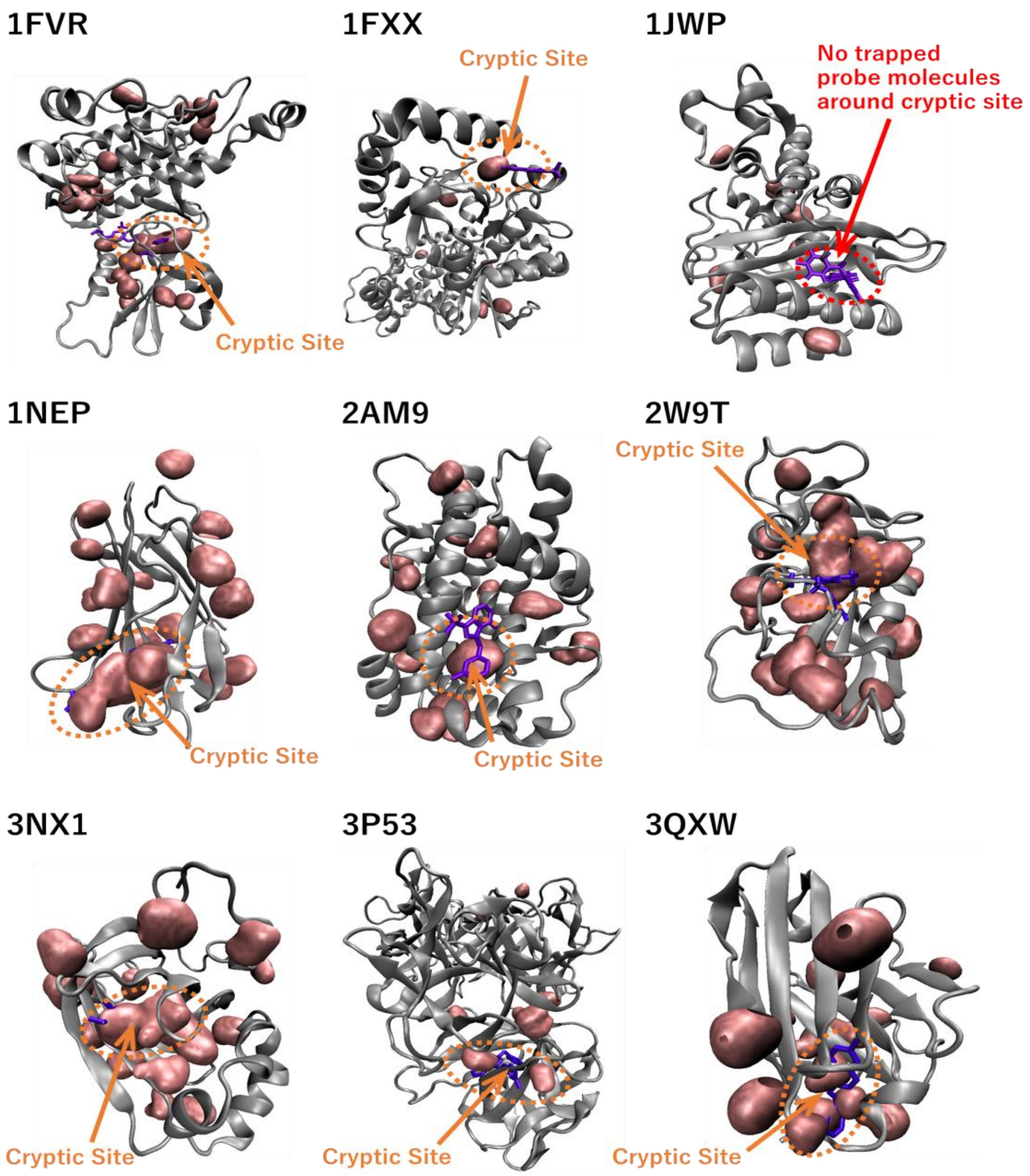


**Figure S2.** Hotspot based on accumulation of probe molecules that could be candidate regions for cryptic sites, obtained by mixed-solvent molecular dynamics simulations. Purple bars and pink regions indicate binding ligands and candidate hotspots, respectively.

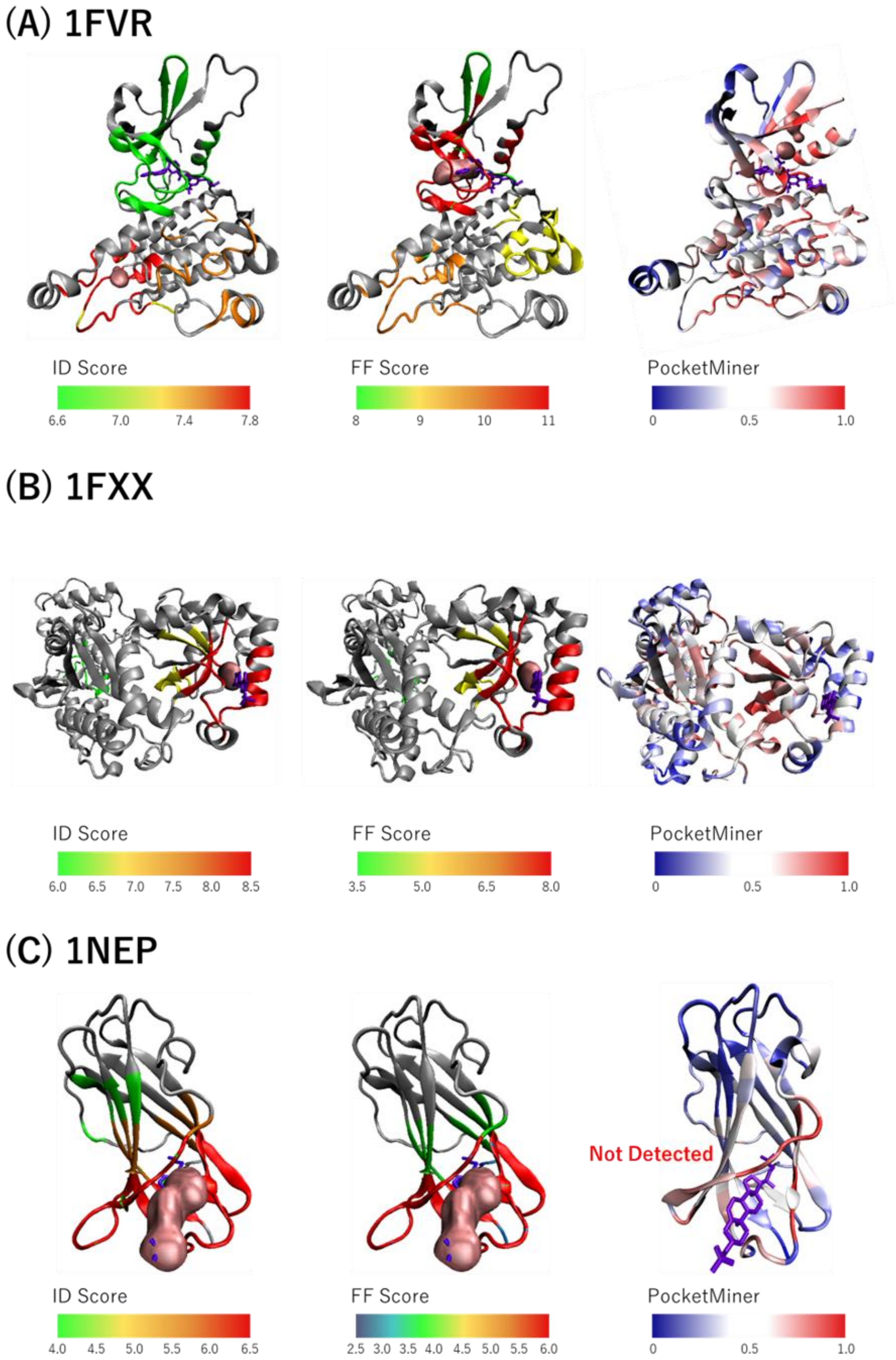
(A) 1FVR
ID Score
6.6
7.0
7.4
7.8
FF Score
8
9
10
11
PocketMiner
0
0.5
1.0
(B) 1FXX
ID Score
6.0
6.5
7.0
7.5
8.0
8.5
FF Score
3.5
5.0
6.5
8.0
PocketMiner
0
0.5
1.0
(C) 1NEP
Not Detected
ID Score
4.0
4.5
5.0
5.5
6.0
6.5
FF Score
2.5
3.0
3.5
4.0
4.5
5.0
5.5
6.0
PocketMiner
0
0.5
1.0

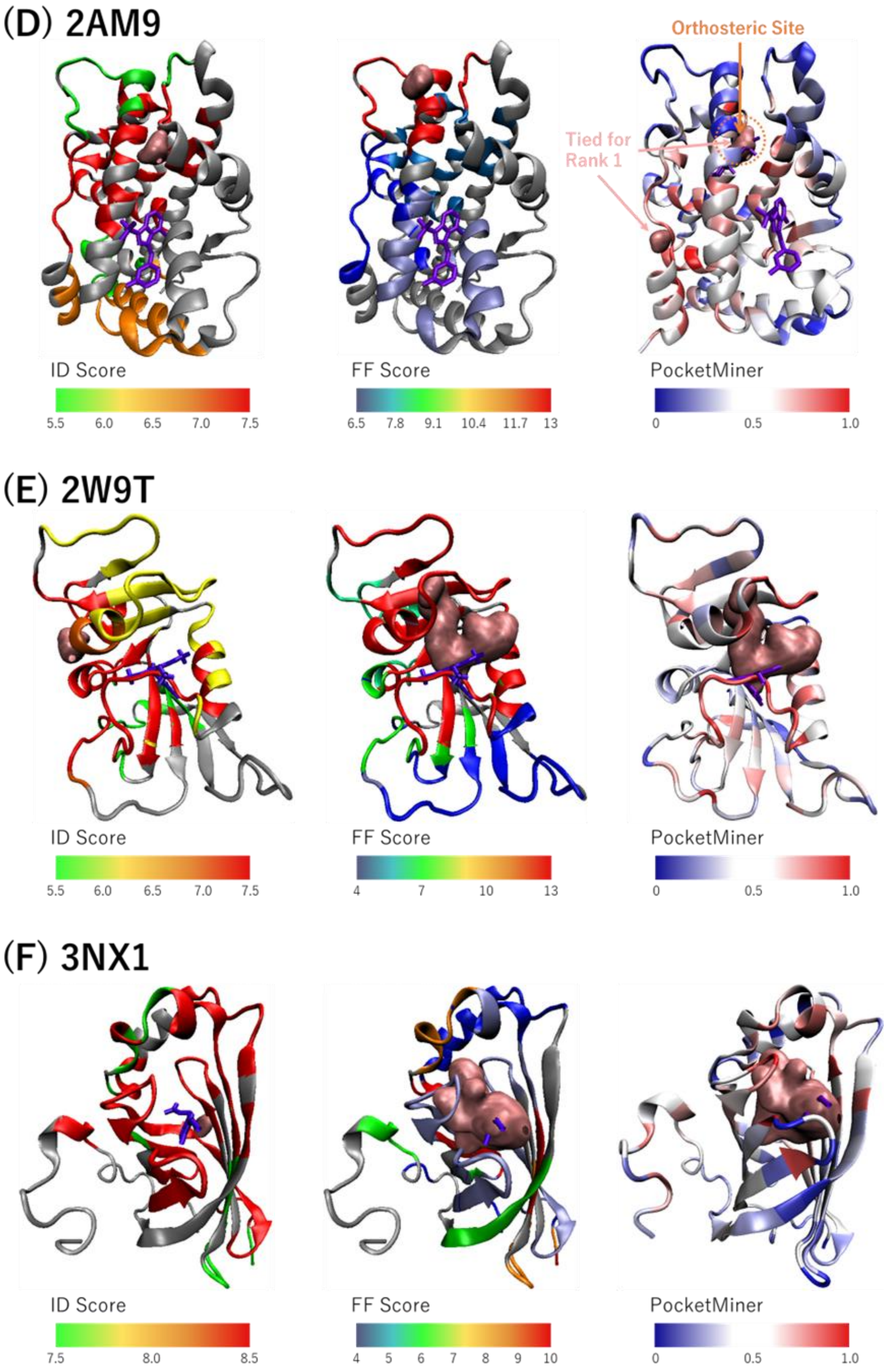

(D) 2AM9
Orthosteric Site
Tied for Rank 1
ID Score
5.5 6.0 6.5 7.0 7.5
FF Score
6.5 7.8 9.1 10.4 11.7 13
PocketMiner
0 0.5 1.0
(E) 2W9T
ID Score
5.5 6.0 6.5 7.0 7.5
FF Score
4 7 10 13
PocketMiner
0 0.5 1.0
(F) 3NX1
ID Score
7.5 8.0 8.5
FF Score
4 5 6 7 8 9 10
PocketMiner
0 0.5 1.0

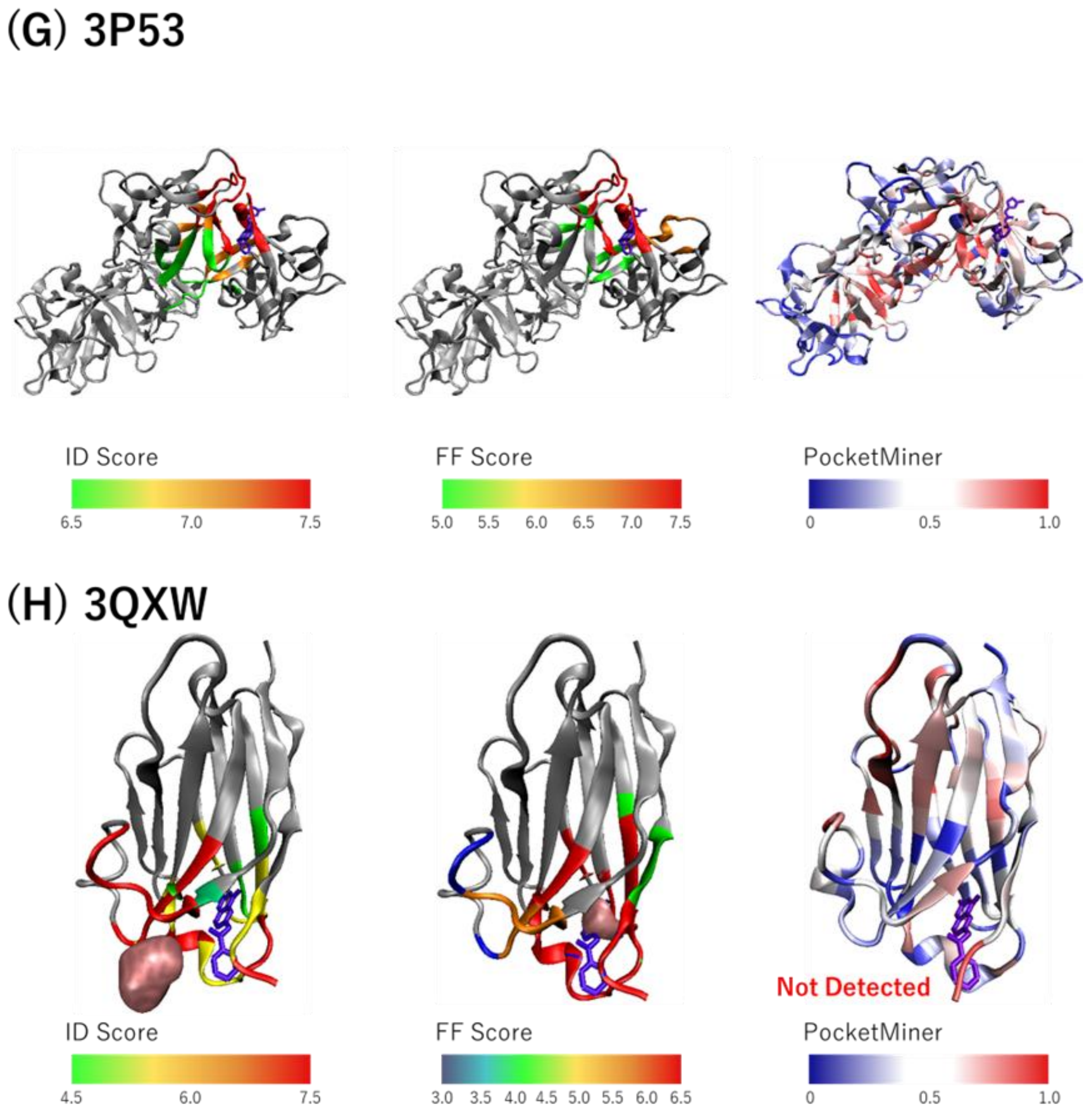


**Figure S3.** The projection views to compare three scores (absolute ID score, FF score, and PocketMiner score) that indicate the cryptic site possibility in the target protein. In these figures of ID and FF scores, the DAIS method was applied using trajectories in phenol-mixed aqueous solvents, after sufficient equilibration of the Phenol mixtures by molecular dynamics. Only the top five scores of ID and FF scores were mapped on protein structures by the color gradations.

**Supplementary Table**

**Table S1. Detection of the residues crashing with the ligands in Exodeoxyribonuclease I**

| List of representative HOLO and APO structures used for detection of clash residues | |
|---|---|
| PDB ID of representative HOLO structures (ligand ID) | PDB ID of representative APO structures |
| 3HL8 (BBP), 3HP9 (CF1) | 1FXX, 2QFX, 4HCB, 4JS5 |

Ratio of clash at residue level in representative APO structures

| Residue | Ligand | |
|---|---|---|
| | BBP | CF1 |
| LYS227 | 0.00 | 0.25 |
| PHE228 | 0.50 | 0.00 |
| TRP245 | 1.00 | 1.00 |
| LEU264 | 0.75 | 0.00 |
| ALA312 | 0.75 | 1.00 |
| ASN313 | 0.25 | 0.75 |
| ARG327 | 0.75 | 1.00 |
| CYS330 | 0.00 | 1.00 |
| LEU331 | 0.25 | 1.00 |
| ARG338 | 0.00 | 1.00 |

**Table S2. Detection of the residues clashing with the ligands in Beta-lactamase TEM**

List of representative HOLO and APO structures used for detection of crash residues

| PDB ID of representative HOLO structures (ligand ID) | PDB ID of representative APO structures |
|---|---|
| 1PZO (CBT), 1PZP (FTA) | 1AXB, 1NYM, 4QY5, 4R4R, 7QLPM, 8DE2 |

Ratio of clash at residue level in representative APO structures

| Residue | Ratio of clash in six representative apo forms | |
|---|---|---|
| | CBT | FTA |
| ALA215 | 0.50 | 0.17 |
| GLY216 | 0.67 | 0.50 |
| PRO217 | 0.83 | 0.50 |
| LEU218 | 1.00 | 0.83 |
| LEU219 | 1.00 | 0.83 |
| ARG220 | 0.67 | 0.17 |
| SER221 | 0.83 | 0.50 |
| ALA222 | 1.00 | 0.83 |
| SER233 | 0.00 | 0.67 |
| GLY234 | 0.00 | 0.17 |
| ALA235 | 0.00 | 0.33 |
| SER240 | 0.00 | 0.33 |
| ARG241 | 0.50 | 0.67 |
| GLY242 | 0.00 | 0.50 |
| ILE243 | 0.00 | 0.17 |
| ARG255 | 0.17 | 0.17 |
| VAL257 | 0.33 | 0.67 |
| ILE259 | 0.33 | 0.33 |

| | | |
|---|---|---|
| GLU270 | 0.33 | 0.33 |
| ASN272 | 0.67 | 0.33 |
| ARG273 | 0.33 | 0.33 |
| GLN274 | 0.33 | 0.33 |
| ILE275 | 0.17 | 0.33 |
| ALA276 | 0.67 | 0.67 |
| GLU277 | 0.00 | 0.33 |
| GLY279 | 0.17 | 0.67 |
| ALA280 | 0.00 | 0.33 |

**Table S3. Detection of the residues clashing with the ligands in Androgen receptor**

| List of representative HOLO and APO structures used for detection of crash residues | |
|---|---|
| PDB ID of representative HOLO structures (ligand ID) | PDB ID of representative APO structures |
| 2PIP (K10), 2PIW (T3), 2YLP (O56) | 1GS4, 1XQ3, 2PNU., 3B5R, 7ZTX |

Ratio of clash at residue level in representative APO structures

| Residue | Ligand | | |
|---|---|---|---|
| | K10 | O56 | T3 |
| VAL717 | 0.80 | 0.40 | 0.00 |
| LYS721 | 0.80 | 0.80 | 0.00 |
| ALA722 | 0.20 | 0.00 | 0.00 |
| VAL731 | 0.40 | 1.00 | 0.00 |
| GLN734 | 0.00 | 0.60 | 0.00 |
| MET35 | 0.80 | 1.00 | 0.00 |
| ALA736 | 0.20 | 0.20 | 0.00 |
| ILE738 | 1.00 | 0.00 | 0.00 |
| GLN739 | 1.00 | 0.80 | 0.00 |
| MET895 | 0.20 | 0.40 | 0.00 |
| MET896 | 0.00 | 0.20 | 0.00 |
| ILE900 | 0.20 | 0.00 | 0.00 |

**Table S4. Detection of the residues crashing with the ligands in Ferulic acid decarboxylase**

List of representative HOLO and APO structures used for detection of crash residues

| PDB ID of representative HOLO structures (ligand ID) | PDB ID of representative APO structures |
|---|---|
| 3NX2 (FER) | 3NX1, 4UU2, 4UU3 |

Ratio of crash at residue level in representative APO structures

| Residue | Lignad |
|---|---|
| | FER |
| TYR21 | 1.00 |
| ASN23 | 1.00 |
| GLY24 | 1.00 |
| TRP25 | 1.00 |
| TYR27 | 1.00 |
| VAL46 | 0.67 |
| PHE95 | 1.00 |
| ILE132 | 1.00 |

**Table S5. Detection of the residues crashing with the ligands in Fascin**

List of representative HOLO and APO structures used for detection of crash residues

| PDB ID of representative HOLO structures (ligand ID) | PDB ID of representative APO structures |
|---|---|
| 6B0T(C7Z), 6I0Z (GZQ), 6I15 (GZT), 6I16 (H0B), 6I17 (GZW), 6I18 (H0N) | 1DFC, 4GP3, 3P53 |

Ratio of crash at residue level in representative APO structures

| Residue | Ligands | | | | | |
|---|---|---|---|---|---|---|
| | C7V | GZQ | GZT | GZW | H0B | H0N |
| PHE14 | 0.00 | 1.00 | 0.00 | 0.33 | 0.00 | 0.33 |
| LEU16 | 0.67 | 1.00 | 0.67 | 0.33 | 0.00 | 0.33 |
| GLN50 | 0.33 | 0.00 | 0.00 | 0.00 | 0.00 | 0.00 |
| SER57 | 0.00 | 0.00 | 0.00 | 0.67 | 0.00 | 0.00 |
| ALA58 | 1.00 | 0.00 | 0.67 | 1.00 | 1.00 | 1.00 |
| ALA59 | 1.00 | 0.00 | 0.00 | 1.00 | 0.00 | 0.00 |
| VAL60 | 1.00 | 0.00 | 1.00 | 0.00 | 0.00 | 0.00 |
| LEU92 | 0.00 | 0.00 | 0.00 | 0.67 | 0.00 | 1.00 |
| ILE93 | 1.00 | 0.00 | 1.00 | 1.00 | 0.67 | 1.00 |
| VAL94 | 0.00 | 0.00 | 0.00 | 0.67 | 1.00 | 1.00 |
| ALA95 | 0.00 | 0.00 | 0.00 | 1.00 | 1.00 | 1.00 |
| TRP101 | 1.00 | 1.00 | 1.00 | 1.00 | 1.00 | 1.00 |
| SER102 | 0.33 | 0.00 | 0.67 | 0.67 | 0.67 | 0.67 |
| LEU103 | 1.00 | 0.00 | 1.00 | 1.00 | 1.00 | 1.00 |
| VAL134 | - | 1.00 | - | - | - | - |
| ILE136 | 0.00 | 0.67 | 0.00 | 0.00 | 0.00 | 0.00 |
| THR213 | 0.00 | 1.00 | 0.00 | 0.00 | 0.00 | 0.00 |
| LEU214 | 1.00 | 1.00 | 1.00 | 1.00 | 1.00 | 1.00 |
| GLU215 | 1.00 | 1.00 | 1.00 | 1.00 | 1.00 | 1.00 |

| | | | | | | |
|---|---|---|---|---|---|---|
| PHE216 | 1.00 | 0.67 | 1.00 | 1.00 | 1.00 | 1.00 |
| ARG217 | 1.00 | 0.00 | 1.00 | 1.00 | 1.00 | 1.00 |
| SER218 | 0.67 | 0.00 | 0.67 | 0.00 | 0.33 | 0.00 |
| GLY219 | 0.67 | 0.00 | 0.00 | 0.00 | 0.00 | 0.00 |
| ARG224 | 0.00 | 0.00 | 0.00 | 1.00 | 0.00 | 0.33 |
| TYR230 | 0.00 | 0.00 | 0.00 | 1.00 | 0.00 | 0.67 |

**Table S6.** Rank order of ID scores calculated by analysis using the DAIS method on the sampling structure after removal of each search molecule and rearrangement of water molecules. If a cryptic site straddled multiple spots, the top rank was selected.

| | Benzene | Isopropanol | Phenol | Imidazole | Acetonitrile | Ethylene glycol | Number of Hotspot |
|---|---|---|---|---|---|---|---|
| 1FVR | 11 | 8 | 4 | 8 | 1 | 9 | 21 |
| 1FXX | 6 | 5 | 1 | 4 | 2 | 5 | 6 |
| 1JWP | --- No trapped probe molecules around the cryptic site --- | | | | | | |
| 1NEP | 4 | 5 | 1 | 1 | 2 | 1 | 16 |
| 2AM9 | 3 | 5 | 12 | 11 | 9 | 5 | 18 |
| 2W9T | 5 | 3 | 2 | 1 | 2 | 1 | 16 |
| 3NX1 | 1 | 1 | 5 | 3 | 3 | 2 | 16 |
| 3P53 | 3 | 3 | 1 | 5 | 3 | 5 | 7 |
| 3QXW | 1 | 1 | 2 | 1 | 1 | 1 | 12 |

**Table S7.** Rank order of FF score calculated by analysis using the DAIS method on the sampling structure after removal of each search molecule and rearrangement of water molecules. If a cryptic site straddled multiple spots, the top rank was selected. The number in parentheses is the total number of candidate sites.

| | Benzene | Isopropanol | Phenol | Imidazole | Acetonitrile | Ethylene glycol | Number of Hotspots |
|---|---|---|---|---|---|---|---|
| 1FVR | 5 | 2 | 1 | 1 | 3 | 5 | 21 |
| 1FXX | 6 | 3 | 1 | 5 | 2 | 2 | 6 |
| 1JWP | --- No trapped probe molecules around the cryptic site --- | | | | | | |
| 1NEP | 1 | 2 | 1 | 1 | 3 | 4 | 16 |
| 2AM9 | 11 | 5 | 2 | 8 | 3 | 3 | 18 |
| 2W9T | 6 | 2 | 1 | 1 | 1 | 1 | 16 |
| 3NX1 | 1 | 1 | 1 | 1 | 1 | 1 | 16 |
| 3P53 | 1 | 1 | 1 | 1 | 2 | 3 | 7 |
| 3QXW | 1 | 2 | 1 | 1 | 2 | 1 | 12 |
